# Prediction of Gene Regulatory Connections with Joint Single-Cell Foundation Models and Graph-Based Learning

**DOI:** 10.1101/2024.12.16.628715

**Authors:** Sindhura Kommu, Yizhi Wang, Yue Wang, Xuan Wang

## Abstract

**Motivation:** Single-cell RNA sequencing (scRNA-seq) data offers unprecedented opportunities to infer gene regulatory networks (GRNs) at a fine-grained resolution, shedding light on cellular phenotypes at the molecular level. However, the high sparsity, noise, and dropout events inherent in scRNA-seq data pose significant challenges for accurate and reliable GRN inference. The rapid growth in experimentally validated transcription factor-DNA binding data (e.g., ChIP-seq) has enabled supervised machine learning methods, which rely on known gene regulatory interactions to learn patterns, and achieve high accuracy in GRN inference by framing it as a gene regulatory link prediction task. This study addresses the gene regulatory link prediction problem by learning informative vectorized representations at the gene level to predict missing regulatory interactions. However, a higher performance of supervised learning methods requires a large amount of known TF-DNA binding data, which is often experimentally expensive and therefore limited in amount. Advances in large-scale pre-training and transfer learning provide a transformative opportunity to address this challenge. In this study, we leverage large-scale pre-trained models, trained on extensive scRNA-seq datasets and known as single-cell foundation models (scFMs). These models are combined with joint graph-based learning to establish a robust foundation for gene regulatory link prediction.

**Results:** We propose scRegNet, a novel and effective framework that leverages scFMs with joint graph-based learning for gene regulatory link prediction. scRegNet achieves state-of-the-art results in comparison with nine baseline methods on seven scRNA-seq benchmark datasets. In addition, scRegNet is more robust than the baseline methods on noisy training data.

**Availability:** The source code is available at https://github.com/sindhura-cs/scRegNet.

## 1. Introduction

Single-cell RNA sequencing (scRNA-seq) has revolutionized the study of cellular diversity by enabling gene expression profiling at an unprecedented single-cell resolution (Jovic et al., 2022). This revolutionary technology has opened up new possibilities for understanding cellular identity, function, and heterogeneity (Linnarsson and Teichmann, 2016). One of the most critical applications of scRNA-seq is the inference of gene regulatory networks (GRNs), which represent the intricate interactions between transcription factors (TFs) and their target genes (Cramer, 2019). These networks govern cellular processes such as differentiation, development, and response to environmental stimuli, making GRN inference crucial for deciphering the molecular mechanisms that bridge genotypes to phenotypes (Alon, 2007). However, accurate prediction of gene regulatory interactions in the GRN poses significant challenges. scRNA-seq data are typically sparse due to high dropout rates (Kharchenko et al., 2014), which occur when some gene transcripts are not captured during sequencing.

Numerous computational methods have been developed to infer gene regulatory interactions in a GRN from scRNA-seq data. Unsupervised approaches, such as GENIE3 (Huynh-Thu et al., 2010) and GRNBoost2 (Moerman et al., 2019), employ tree-based regression techniques to identify gene sets co-expressed with transcription factors (TFs). However, these methods face notable challenges. Given the large number of genes profiled and the relatively smaller number of samples, many co-expression or co-functionality signals may arise purely from chance or noise in the data (Freytag et al., 2015). Recently, supervised learning-based methods have gained attention by using experimentally validated TF-DNA binding data from resources such as ENCODE (Consortium, 2012), ChIP-Atlas (Okanishi et al., 2021), and ESCAPE (Xu et al., 2015) to train deep learning models.

The supervised deep learning models have achieved significantly higher accuracy than unsupervised methods by learning from known TF-gene pairs to predict missing gene regulatory interactions. One such supervised method, CNNC (Yuan and Bar-Joseph, 2019), employs convolutional neural networks (CNNs) for gene regulatory link prediction by converting gene pair co-expression profiles into image-like histograms. Another supervised method, GNE (Kc et al., 2019), uses multilayer perceptrons (MLPs) to encode gene expression profiles and graph topologies for gene regulatory link prediction. More recently, graph-based learning frameworks, such as GENELink (Chen and Liu, 2022) and GNNLink (Mao et al., 2023), have demonstrated promise in modeling the complex interconnections within GRNs with graph neural networks (GNNs). However, a higher performance of supervised learning methods requires a large amount of experimentally validated gene regulatory networks as labeled training data, which is often experimentally expensive and therefore limited in amount.

Advances in large-scale pre-training and transfer learning offer a transformative opportunity to address the above challenge. Large-scale pre-trained foundation models have significantly impacted fields such as natural language understanding and computer vision by utilizing deep learning models pre-trained on large-scale text or image datasets, which can then be used for various downstream tasks with limited task-specific data. Similarly, for scRNA-seq data, large-scale pre-trained foundation models have become essential tools for interpreting the “languages” of cells. Models such as scFoundation (Hao et al., 2024), Geneformer (Theodoris et al., 2023), scBERT (Yang et al., 2022), and scGPT (Cui et al., 2024) leverage extensive unlabeled scRNA-seq datasets to learn context-aware representations of genes in a single-cell, capturing latent gene-gene interactions across the genome. These models, known as single-cell foundation models (scFMs), trained on large-scale scRNA-seq data spanning millions of samples, provide rich informative representations for advancing network biology.

In this study, we propose a novel and effective framework, scRegNet, that combines scFMs with joint graph-based learning for gene regulatory link prediction. scRegNet harnesses the rich, context-aware gene-level representations learned by large-scale pre-trained models and combines them with gene-level representations derived from graph-based learning for predicting gene regulatory interactions. This integration enables the model to leverage both the contextual gene interaction patterns (learned from the whole genome expression of millions of unlabelled scRNA-seq data) and regulatory network topology (learned from graph-based encoders), essential for accurate gene regulatory link prediction. We evaluated scRegNet on seven single-cell scRNA-seq benchmark datasets from BEELINE (Pratapa et al., 2020), covering a diverse range of cell types from both human and mouse sources. scRegNet consistently surpasses nine state-of-the-art baseline methods, demonstrating significant improvements in both the Area Under the Receiver Operating Characteristic Curve (AUROC) and the Area Under the Precision-Recall Curve (AUPRC) across all seven benchmark datasets. In addition, our experiments demonstrate that scRegNet is more robust compared to the baseline methods on noisy training data.

## 2. Method

In this section, we discuss each element of scRegNet (Fig 1) in detail.

**Fig. 1:**
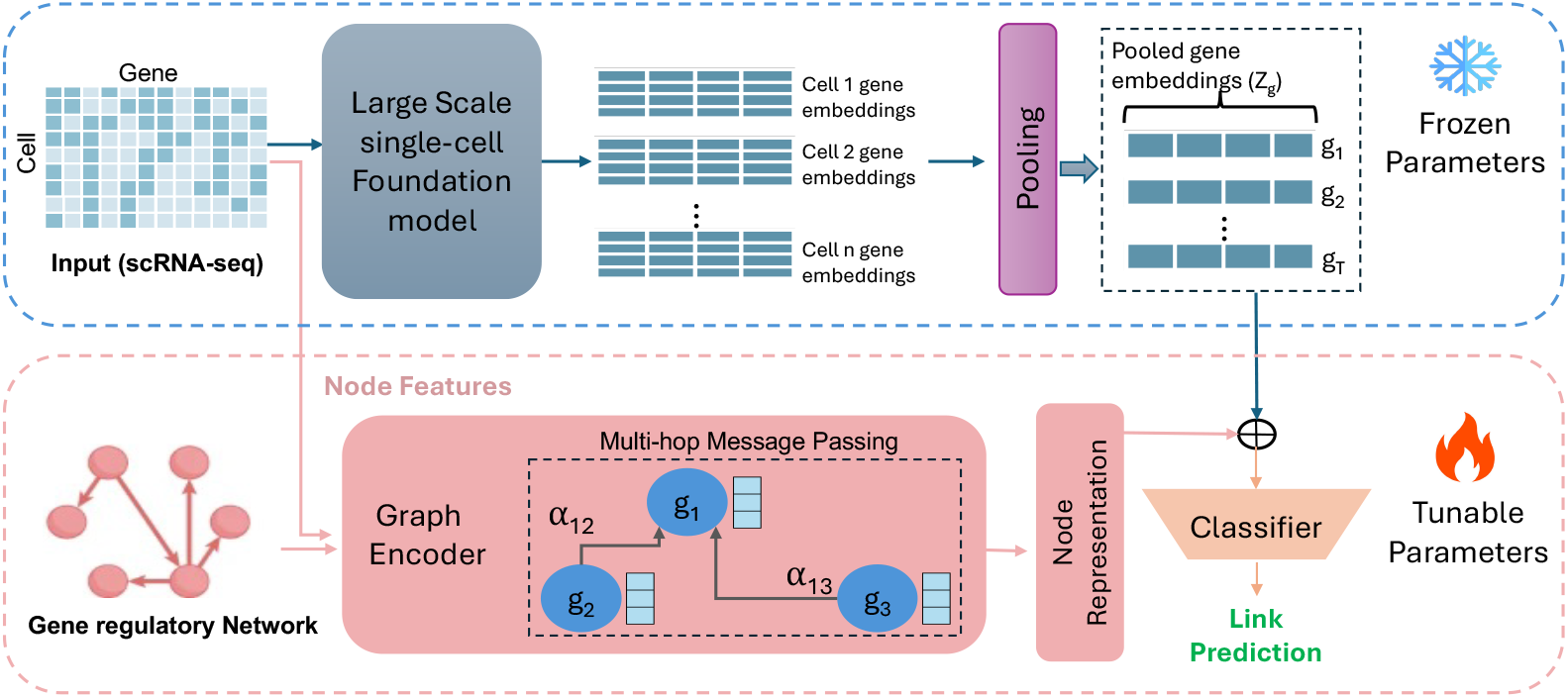
Overview of the scRegNet framework for GRN inference. scRegNet utilizes a pre-trained single-cell foundation model (top; Section 2.1) to generate gene embeddings from scRNA-seq input which are integrated with outputs from a Graph Encoder (bottom left; Section 2.2). The combined representations are fed into a classifier (bottom right; Section 2.4) for link prediction, enabling the identification of missing regulatory interactions among genes. The architecture incorporates both frozen parameters for leveraging pre-trained knowledge and tunable parameters for domain-specific learning, facilitating the seamless integration of biological context with learned embeddings.

### 2.1. Gene representations from foundation models

Recent studies (Hao et al., 2024; Theodoris et al., 2023; Yang et al., 2022; Cui et al., 2024), have demonstrated that large-scale pre-trained foundation models possess a strong capacity to model gene-gene interactions across cells, achieving state-of-the-art performance in various single-cell analysis tasks. In this study, we explore three single-cell foundation models (scFMs), scBERT (Yang et al., 2022), Geneformer (Theodoris et al., 2023), and scFoundation (Hao et al., 2024), to capture the context-aware gene-gene relationships of the scRNA-seq data. A summary of these three scFMs, including their architectures and key features, is provided in Supplementary Table S1. All of these three scFMs rely on attention-based Transformer architectures (Vaswani et al., 2017) for processing gene-level vector representations of the scRNA-seq data and employ masked language modeling (MLM) as a self-supervised pre-training strategy to learn multifaceted internal patterns of cells from millions of single-cell transcriptomes. The MLM strategy is the same as that used in pre-training the large language models (LLMs), such as ChatGPT, allowing the LLMs to learn human knowledge from huge archives of natural language texts. However, these three scFMs differ in how they represent the input scRNA-seq data, their model architectures, and their training procedures. Specifically, the input design and pre-processing steps vary for each model as detailed below.

First, we formally define the input scRNA-seq data as a cell-by-gene matrix, **X** ∈ R^*N ×T*^, where each element represents the RNA abundance for gene *t* in cell *n*. This matrix, referred to as the raw count matrix, is normalized using a log transformation and feature scaling to ensure compatibility with attention-based architectures. To create a sequence suitable for as input for these models, we define a sequence of gene tokens as {*g*_1_, …, *g*_*T*_}, where *T* is the total number of selected genes in the dataset. Then we go into details of each scFM in how they handle this input data.

#### 2.1.1. scBERT

scBERT (Yang et al., 2022) utilizes a combination of two features for each gene: (1) a gene ID feature with gene2vec (Du et al., 2019) that represents individual genes in a pre-defined vector space, and (2) a gene expression level feature. For each gene token *g*_*t*_, the initial input representation is constructed as 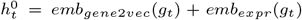, where emb_gene2vec_(·) denotes the gene identity embedding and emb_expr_(·) represents the expression level embedding. These input representations are processed through *L* = 6 successive transformer encoder layers:

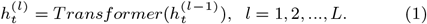

The final hidden states 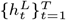 serve as the 200-dimensional gene-level embeddings, suitable for downstream tasks. See Supplementary S1.1 for further details.

#### 2.1.2. scFoundation

scFoundation (Hao et al., 2024) utilizes an asymmetric encoder-decoder architecture (Gong et al., 2024) that employs attention mechanisms to optimize gene dependency extraction in sparse single-cell data. It also includes an embedding module that converts continuous gene expression scalars into high-dimensional vectors, allowing the model to fully retain the information from raw expression values, rather than discretizing them like other methods. The encoder is designed to only process non-zero and non-masked gene expression embeddings. These encoded embeddings are then recombined with the zero-expressed gene embeddings at the decoder stage to produce final 512-dimensional gene-level representations. These vector representations capture detailed gene dependencies, making them suitable for downstream network biology-based tasks. See Supplementary S1.2 for further details.

#### 2.1.3. Geneformer

Geneformer (Theodoris et al., 2023) employs a rank value encoding strategy to represent input scRNA-seq data, prioritizing genes based on their expression value within a cell. To prepare the input data for Geneformer, we utilize the token dictionary (TokenDict) and the gene median file (GeneMedian) provided in the model’s repository. These resources ensure that the input is accurately tokenized based on the rank value encoding strategy, maintaining consistency with the pre-trained model. Each gene’s expression is normalized relative to a median reference and then converted into ranked tokens. The input matrix **R** is constructed as follows:

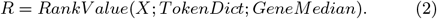

The RankValue function normalizes each gene’s expression using the GeneMedian values and maps them to discrete tokens via the TokenDict. For each single-cell transcriptome, Geneformer embeds each gene into an 896-dimensional space that captures the gene’s contextual characteristics within the cell. These contextual embeddings are generated via multi-layer attention mechanisms similar to the equation 1, where 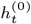 is the initial embedding for token *R*_*t*_, and *L* = 20 layers are used in the encoder. The final hidden state, 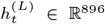, represents the context-aware embedding for gene *t*. To obtain robust and generalizable gene representations, embeddings are extracted from the penultimate layer of the model, as it captures a more abstract and general feature space compared to the final layer. See Supplementary S1.3 for further details.

#### 2.1.4. Mean pooling

Upon extracting the gene representations as gene embeddings from the three scFMs as described above, the gene embeddings for all the cells under each cell type are further aggregated together at the gene level to establish a cohesive representation for each gene. This is accomplished through mean pooling. The mean-pooled embedding for gene *t* is computed as follows:

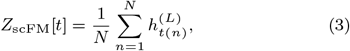

where 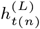 represents the extracted embedding of gene *t* within cell *n* (sections 2.1.1-2.1.3). This pooling methodology facilitates a balanced representation that encapsulates the average gene activity throughout the dataset.

### 2.2. Graph-based learning with GNNs

In addition to the context-aware gene-level representations extracted from the scFMs mentioned above, we also extract gene representations that encode the regulatory network topology using GNNs. The regulatory network topology comes from the gene interactions in the training data using experimentally validated TF-DNA binding data from resources such as ENCODE (Consortium, 2012), ChIP-Atlas (Okanishi et al., 2021), and ESCAPE (Xu et al., 2015). We formulate these gene interactions in the training data as a known graph between TFs and target genes, with nodes denoting TFs or genes and links symbolizing their regulatory associations. The gene regulatory link prediction task aims to discover any missing interactions between gene pairs that are not included in the training data. Specifically, given the gene interactions in the training data, the graph encoders (GNNs) learn a mapping function that can generate low-dimensional gene embeddings that capture the underlying structure of the gene interactions.

Let the gene interactions in the training data be represented as *G* = {*V, E*}, where *V* is the set of nodes (genes) and E is the set of edges (regulatory interactions). The goal is to learn effective node representations through message passing, which embeds into each node—information about its multi-hop neighbors. Specifically, each node receives and aggregates messages (i.e., features or embeddings) from its neighboring nodes recursively in multiple layers Formally, the updated representation *v*_*t*_^*l*^ of each node, in *l*-th layer is given by:

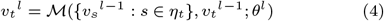

where *η*_*t*_ represents the set of neighboring nodes for an arbitrary node *t*, and ℳ (·), parameterized by *θ*_*l*_ in the *l*-th layer, is the message passing function for neighborhood aggregation. The neighborhood aggregation varies depending on the type of GNN.

To derive the initial features of the genes, we apply pre-processing operations to the raw single-cell expression data. Here in scRegNet, a simple Graph Convolutional Network (GCN) (Kipf and Welling, 2017) is employed. We noticed that this simple architecture is adequate to reach similar performance compared to computation-demanding GNN frameworks such as Graph Attention Networks (GAT) (Veličković et al., 2018) and GraphSAGE (SAmple and aggreGatE) (Hamilton et al., 2017). A detailed GNN framework comparison and analysis can be found in the Results section.

### 2.3. Unified gene representations

After extracting gene representations from both the scFMs and the GNNs, we integrate them into a unified gene representation for each gene as shown in Fig 1. This integration involves concatenating the representations from scFM (capturing contextual gene interactions) and GNN (capturing network topology) as follows:

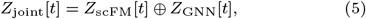

where *Z*_scFM_[*t*] represents the foundation model-derived representation for gene *t* (section 2.1) and *Z*_GNN_[*t*] represents the node representation of gene *t* derived from the GNN encoder (section 2.2). The concatenated representation *Z*_joint_[*t*] serves as the unified representation for gene *t*.

### 2.4. Link prediction layer

The link prediction module constitutes the final component of scRegNet, specifically designed to evaluate the likelihood of unseen regulatory interactions between the gene pairs. We employ Multilayer Perceptron (MLP) networks integrated with ReLU activation functions and Dropout regularization for this task.

For each gene pair (*i, j*), unified feature representations *Z*_joint_[*i*] and *Z*_joint_[*j*] as derived above are processed through MLP. The two outputs from the MLP are concatenated to form a combined representation that captures the joint features of the gene pair. This concatenated representation is passed to a fully connected classification layer. This classification layer predicts the likelihood of a regulatory interaction by outputting a score for each possible class (presence or absence of an interaction). The predicted scores are normalized using a softmax function, yielding probabilities for each class, as shown in the following equation:

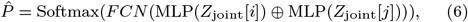

where ⊕ represents the concatenation operation, and *FCN* denotes the final fully connected network that maps the combined representation to the output probabilities. The predicted probabilities correspond to the likelihood of the presence (*Ŷ* = 1) or absence (*Ŷ* = 0) of a regulatory interaction.

### 2.5. Model training

To train the scRegNet model, we employ the Binary Cross-Entropy (BCE) loss function, which measures the difference between the predicted regulatory interaction probabilities and the ground-truth labels in the training dataset:

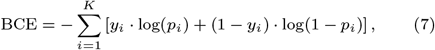

where *K* is the total number of gene pairs in training data, *y*_*i*_ is the ground-truth label for the *i*-th gene pair (*y*_*i*_ = 1 for interaction, *y*_*i*_ = 0 for no interaction), *p*_*i*_ is the predicted probability of a regulatory interaction for the *i*-th pair. The BCE loss is backpropagated through the scRegNet framework, enabling end-to-end optimization of the model parameters. The parameters of GNN layers are updated while the parameters of the scFM remain frozen during training as shown in Fig 1. Refer to the Supplementary S4 for a comprehensive step-by-step training algorithm.

## 3. Experimental Setup

### 3.1. Datasets and data pre-processing

We evaluate scRegNet on seven scRNA-seq benchmark datasets provided in BEELINE (Pratapa et al., 2020), more specifically 1) human embryonic stem cells (hESC), 2) human mature hepatocytes (hHEP), 3) mouse dendritic cells (mDC), 4) mouse embryonic stem cells (mESC), 5) mouse hematopoietic stem cells of the erythroid lineage (mHSC-E), 6) mouse hematopoietic stem cells with a granulocyte-monocyte lineage (mHSC-GM), and 7) mouse hematopoietic stem cells with a lymphoid-like lineage (mHSC-L). Following GENELink (Chen and Liu, 2022) and GNNLink (Mao et al., 2023), we adopt the cell-type-specific ChIP-seq networks from the aforementioned datasets as ground truth to evaluate the performance of scRegNet and baseline methods.

Following the original paper of BEELINE (Pratapa et al., 2020) that provided the seven benchmark datasets, we pre-process each scRNA-seq dataset by only inferring the interactions outgoing from TFs. Following BEELINE (Pratapa et al., 2020), we respectively select 500 and 1000 significantly most-varying genes with all TFs whose corrected P-value (Bonferroni method) of variance is lower than 0.01 as the ground truth network for gene regulatory link prediction. The seven scRNA-seq datasets can be downloaded from Gene Expression Omnibus with the accession numbers GSE81252 (hHEP), GSE75748 (hESC), GSE98664 (mESC), GSE48968 (mDC) and GSE81682 (mHSC).

For a fair comparison with existing state-of-the-art baseline models (Section 3.3), we follow the same evaluation strategy as GENELink (Chen and Liu, 2022) to split the ground truth networks into training/validation/test sets in all benchmark datasets. In these ground truth networks, the number of TFs is limited, and most of them are with high degrees. To validate that the supervised model can distinguish the much more subtle differences between target and non-target genes for each TF; we divide the positive and negative target genes of each TF in proportion to the training and test datasets.

Specifically, for each transcription factor (TF), the interactions (edges) with target genes are categorized into positive and negative samples. Positive samples represent true regulatory relationships supported by experimental evidence, such as ChIP-seq data from ground-truth networks. Negative samples, on the other hand, consist of gene pairs with no known regulatory interactions. To ensure a robust evaluation framework, the positive and negative samples for each TF are divided proportionally into training and test sets, maintaining a fixed ratio of 67% for training and 33% for testing. This partitioning ensures a consistent evaluation process across all TFs. Additionally, a small subset of the training data (10%) is reserved as a validation set for hyperparameter tuning and early stopping during model training. The data splitting is performed per TF, ensuring that all TFs contribute examples to both the training and test sets. Crucially, this partitioning strategy prevents data leakage by ensuring that the same gene does not appear in both the training and test sets for the same TF. This approach guarantees that the model’s performance is evaluated on entirely independent data, maintaining the integrity of the evaluation process. The sizes of each ground-truth network training set are listed in Table 1.

**Table 1.**
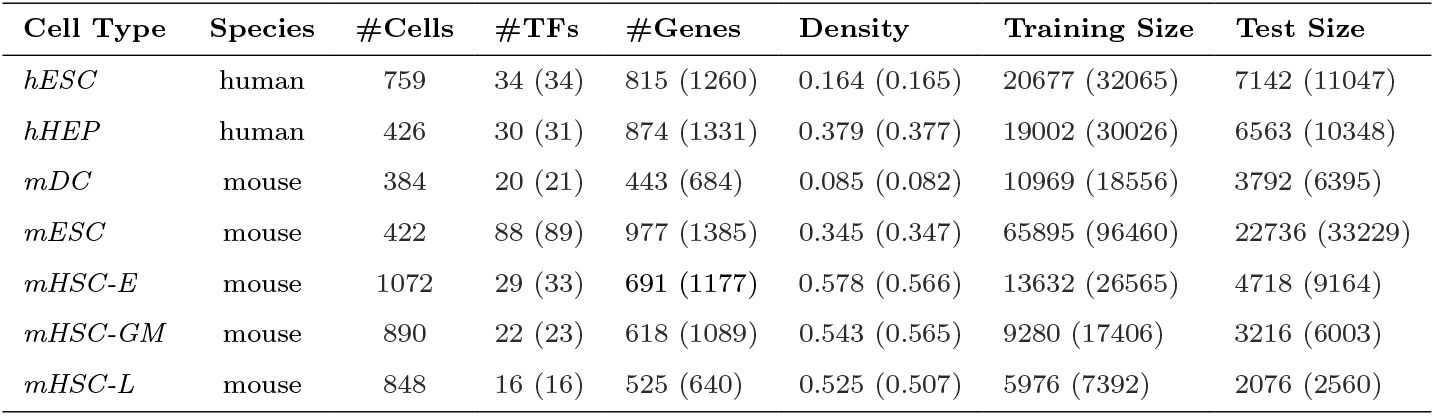
The statistics of prior networks with TFs and 500 (1000) most-varying genes.

### 3.2. Parameter Settings

We leveraged Optuna (Akiba et al., 2019), a powerful hyperparameter optimization framework, to systematically explore the search space. Detailed information on the hyperparameter settings can be found in Supplementary Table S2.

### 3.3. Baseline models and evaluation metrics

We compare scRegNet against nine baseline methods for gene regulatory link prediction from single-cell RNA-seq data, which have been proven to achieve good performance. GNNLink (Mao et al., 2023) and GENELink (Chen and Liu, 2022) utilize GCNs and GATs, respectively, to capture regulatory structures, while GNE (Kc et al., 2019), CNNC (Yuan and Bar-Joseph, 2019), and DeepDRIM (Chen et al., 2021) employ MLPs, CNNs, and supervised learning with gene expression images for prediction. GRN-transformer (Shu et al., 2022) uses axial transformers for weakly supervised inference of cell type-specific GRNs. Traditional methods include PCC (Salleh et al., 2015) for linear correlation, and machine learning models GRNBoost2 (Moerman et al., 2019) and GENIE3 (Huynh-Thu et al., 2010). These models have demonstrated state-of-the-art performance on benchmark datasets, as confirmed by the evaluations in the GENELink (Chen and Liu, 2022) and GNNLink (Mao et al., 2023) papers. See Supplementary S2 for more details on baseline methods. We use the Area Under the Receiver Operating Characteristic Curve (AUROC) and the Area Under the Precision-Recall Curve (AUPRC) as the evaluation metrics.

## 4. Results

### 4.1. Performance on benchmark datasets

As demonstrated in Tables 2 and 3, all variants of scRegNet (w/ scBERT, Geneformer, and scFoundation) consistently surpass existing baseline models in both AUROC and AUPRC metrics across all seven cell-type-specific datasets (hESC, hHEP, mDC, mESC, mHSC-E, mHSC-GM, mHSC-L). The Geneformer- and scFoundation-based configurations achieved the highest performance, with scRegNet(w/ Geneformer) delivering an average improvement of 7.4% and 6.9% in AUROC and 18.6% and 4.1% (Table 2) in AUPRC over GNNLink and GENELink respectively, on datasets with 500 most-variable genes (TFs+500). Similarly, for TFs+1000 datasets, scRegNet(w/ Geneformer) outperformed GENELink and GNNLink by 6.2% and 7.5% in AUROC and 16.5% and 3.9% in AUPRC (Table 3) respectively. Among the three scFM backbone configurations, Geneformer and scFoundation yielded slightly better results compared to scBERT. Traditional methods, such as GRNBOOST2, GENIE3, and PCC, demonstrated limited predictive accuracy, particularly in AUPRC, due to their reliance on simplistic pairwise correlation metrics. In contrast, graph-based deep learning frameworks like GNNLink and GENELink improved performance by leveraging gene-gene interactions, but their effectiveness remained limited. scRegNet consistently outperformed these approaches across diverse cell types and datasets. For example, as shown in Table 2, scRegNet achieved an AUPRC of 0.62 in hESC, representing a +24% improvement over GENELink (AUPRC = 0.50) and a +21.6% improvement over GNNLink (AUPRC = 0.51) under comparable conditions. scRegNet demonstrated superior performance in challenging scenarios with sparse regulatory signals, emphasizing the value of integrating foundation model embeddings to capture context-aware gene relationships and overcome the limitations of correlation-based and graph-only methods.

**Table 2.** Link prediction performance on seven scRNA-seq datasets with **500 most-variable genes**. Each dataset includes a cell-type-specific ground-truth network. The values reported are averages from 50 independent evaluations per cell type. scRegNet utilizing the three backbone models—scBERT, Geneformer, and scFoundation—consistently outperforms the baselines.

| Method |  | Cell Type |  |  |  |  |  |  |
| --- | --- | --- | --- | --- | --- | --- | --- | --- |
|  |  | hESC | hHEP | mDC | mESC | mHSC-E | mHSC-GM | mHSC-L |
| *GRNBOOST2*<br>(Moerman et al., 2019) | AUROC | 0.49 | 0.52 | 0.52 | 0.53 | 0.53 | 0.50 | 0.52 |
|  | AUPRC | 0.15 | 0.38 | 0.06 | 0.32 | 0.57 | 0.52 | 0.5 |
| *GENIE3*<br>(Huynh-Thu et al., 2010) | AUROC | 0.50 | 0.54 | 0.50 | 0.50 | 0.52 | 0.53 | 0.52 |
|  | AUPRC | 0.15 | 0.39 | 0.05 | 0.31 | 0.56 | 0.53 | 0.50 |
| *PCC*<br>(Salleh et al., 2015) | AUROC | 0.47 | 0.49 | 0.54 | 0.51 | 0.49 | 0.54 | 0.55 |
|  | AUPRC | 0.14 | 0.35 | 0.06 | 0.31 | 0.56 | 0.53 | 0.52 |
| *GRN-Transformer*<br>(Shu et al., 2022) | AUROC | 0.51 | 0.49 | 0.50 | 0.53 | 0.64 | 0.50 | 0.64 |
|  | AUPRC | 0.15 | 0.35 | 0.06 | 0.49 | 0.71 | 0.66 | 0.64 |
| *DeepDRIM*<br>(Chen et al., 2021) | AUROC | 0.63 | 0.52 | 0.50 | 0.51 | 0.56 | 0.64 | 0.58 |
|  | AUPRC | 0.13 | 0.39 | 0.06 | 0.46 | 0.76 | 0.64 | 0.59 |
| *CNNC*<br>(Yuan and Bar-Joseph, 2019) | AUROC | 0.68 | 0.64 | 0.54 | 0.73 | 0.67 | 0.69 | 0.67 |
|  | AUPRC | 0.25 | 0.46 | 0.06 | 0.48 | 0.74 | 0.68 | 0.64 |
| *GNE*<br>(Kc et al., 2019) | AUROC | 0.67 | 0.80 | 0.52 | 0.81 | 0.82 | 0.83 | 0.77 |
|  | AUPRC | 0.34 | 0.65 | 0.06 | 0.64 | 0.80 | 0.78 | 0.70 |
| *GENELink*<br>(Chen and Liu, 2022) | AUROC | 0.82 | 0.84 | 0.71 | 0.88 | 0.87 | 0.89 | 0.83 |
|  | AUPRC | 0.50 | 0.70 | 0.11 | 0.75 | 0.89 | 0.89 | 0.83 |
| *GNNLink*<br>(Mao et al., 2023) | AUROC | 0.85 | 0.82 | 0.70 | 0.84 | 0.83 | 0.89 | 0.84 |
|  | AUPRC | 0.52 | 0.75 | **0.25** | 0.76 | 0.88 | 0.89 | 0.85 |
| **scRegNet**<br>(w/ **scBERT**) | AUROC | 0.88±0.00 | **0.90±0.00** | 0.75±0.01 | 0.92±0.00 | **0.92±0.00** | 0.92±0.00 | 0.85±0.01 |
|  | AUPRC | 0.61±0.00 | 0.83±0.00 | 0.12±0.01 | 0.84±0.00 | **0.94±0.00** | 0.93±0.00 | 0.85±0.01 |
| **scRegNet**<br>(w/ **scFoundation**) | AUROC | **0.89±0.00** | **0.90±0.00** | **0.81±0.00** | **0.93±0.00** | **0.92±0.00** | **0.93±0.00** | **0.88±0.00** |
|  | AUPRC | **0.62±0.00** | 0.83±0.00 | 0.15±0.01 | **0.86±0.00** | **0.94±0.00** | **0.94±0.00** | **0.88±0.00** |
| **scRegNet**<br>(w/ **Geneformer**) | AUROC | **0.89±0.00** | **0.90±0.00** | **0.81±0.00** | **0.93±0.00** | **0.92±0.00** | **0.93±0.00** | **0.88±0.00** |
|  | AUPRC | **0.62±0.00** | **0.84±0.00** | 0.17±0.00 | **0.86±0.00** | **0.94±0.00** | **0.94±0.00** | **0.88±0.00** |

**Table 3.** Link prediction performance on seven scRNA-seq datasets with **1000 most-variable genes**. Each dataset includes a cell-type-specific ground-truth network. The values reported are averages from 50 independent evaluations per cell type. scRegNet utilizing the three backbone models—scBERT, Geneformer, and scFoundation—consistently outperforms the baselines.

| Method |  | Cell Type |  |  |  |  |  |  |
| --- | --- | --- | --- | --- | --- | --- | --- | --- |
|  |  | hESC | hHEP | mDC | mESC | mHSC-E | mHSC-GM | mHSC-L |
| *GRNBOOST2*<br>(Moerman et al., 2019) | AUROC | 0.49 | 0.52 | 0.53 | 0.53 | 0.51 | 0.49 | 0.53 |
|  | AUPRC | 0.14 | 0.37 | 0.05 | 0.32 | 0.54 | 0.52 | 0.48 |
| *GENIE3*<br>(Huynh-Thu et al., 2010) | AUROC | 0.50 | 0.54 | 0.52 | 0.50 | 0.50 | 0.51 | 0.52 |
|  | AUPRC | 0.15 | 0.38 | 0.05 | 0.31 | 0.54 | 0.53 | 0.48 |
| *PCC*<br>(Salleh et al., 2015) | AUROC | 0.47 | 0.50 | 0.54 | 0.51 | 0.49 | 0.54 | 0.55 |
|  | AUPRC | 0.14 | 0.34 | 0.05 | 0.31 | 0.53 | 0.54 | 0.51 |
| *GRN-Transformer*<br>(Shu et al., 2022) | AUROC | 0.67 | 0.58 | 0.57 | 0.50 | 0.59 | 0.53 | 0.58 |
|  | AUPRC | 0.16 | 0.53 | 0.05 | 0.51 | 0.69 | 0.61 | 0.52 |
| *DeepDRIM*<br>(Chen et al., 2021) | AUROC | 0.56 | 0.63 | 0.50 | 0.62 | 0.50 | 0.66 | 0.57 |
|  | AUPRC | 0.19 | 0.46 | 0.06 | 0.46 | 0.73 | 0.64 | 0.48 |
| *CNNC*<br>(Yuan and Bar-Joseph, 2019) | AUROC | 0.72 | 0.66 | 0.56 | 0.73 | 0.72 | 0.69 | 0.62 |
|  | AUPRC | 0.27 | 0.49 | 0.05 | 0.50 | 0.77 | 0.73 | 0.56 |
| *GNE*<br>(Kc et al., 2019) | AUROC | 0.68 | 0.81 | 0.52 | 0.82 | 0.84 | 0.84 | 0.77 |
|  | AUPRC | 0.34 | 0.66 | 0.05 | 0.65 | 0.81 | 0.81 | 0.68 |
| *GENELink*<br>(Chen and Liu, 2022) | AUROC | 0.83 | 0.85 | 0.74 | 0.89 | 0.90 | 0.90 | 0.84 |
|  | AUPRC | 0.50 | 0.71 | 0.12 | 0.76 | 0.90 | 0.91 | 0.81 |
| *GNNLink*<br>(Mao et al., 2023) | AUROC | 0.80 | 0.84 | 0.78 | 0.84 | 0.87 | 0.92 | 0.86 |
|  | AUPRC | 0.51 | 0.78 | **0.21** | 0.78 | 0.93 | 0.93 | 0.86 |
| **scRegNet**<br>(w/ **scBERT**) | AUROC | 0.86±0.00 | 0.90±0.00 | 0.75±0.01 | **0.93±0.01** | 0.92±0.00 | 0.92±0.00 | 0.81±0.01 |
|  | AUPRC | 0.56±0.01 | 0.83±0.00 | 0.14±0.01 | 0.86±0.01 | 0.94±0.00 | 0.94±0.00 | 0.79±0.01 |
| **scRegNet**<br>(w/ **scFoundation**) | AUROC | **0.88±0.00** | **0.91±0.00** | 0.82±0.00 | **0.93±0.00** | **0.94±0.00** | **0.94±0.00** | 0.87±0.00 |
|  | AUPRC | **0.62±0.00** | **0.85±0.00** | 0.15±0.00 | 0.86±0.00 | **0.95±0.00** | **0.95±0.00** | 0.86±0.00 |
| **scRegNet**<br>(w/ **Geneformer**) | AUROC | **0.88±0.00** | 0.90±0.00 | **0.84±0.00** | **0.93±0.00** | **0.94±0.00** | **0.94±0.00** | **0.88±0.00** |
|  | AUPRC | **0.62±0.00** | 0.84±0.00 | 0.17±0.01 | **0.87±0.00** | **0.95±0.00** | **0.95±0.00** | **0.87±0.00** |

### 4.2. Ablation Study

To understand the contribution of each component within scRegNet, we performed a series of ablation studies, with the results displayed in Figure 2. The first experiment involved removing the GNN encoder, which led to a substantial decline in performance, highlighting the critical role of graph-based representation learning in refining gene embeddings. In the second ablation, we excluded the pre-trained foundation model embeddings. This omission impaired performance, demonstrating the importance of capturing diverse cellular contexts through pre-trained embeddings for accurate GRN inference. In these experiments, we utilized Geneformer as the scFM backbone and GCN as the GNN backbone for the model. For each dataset, the AUROC score is calculated as the average of the AUROC values from the TF+500 and TF+1000 datasets. Similarly, the AUPRC score is computed as the average of the AUPRC values from these two networks.

**Fig. 2:**
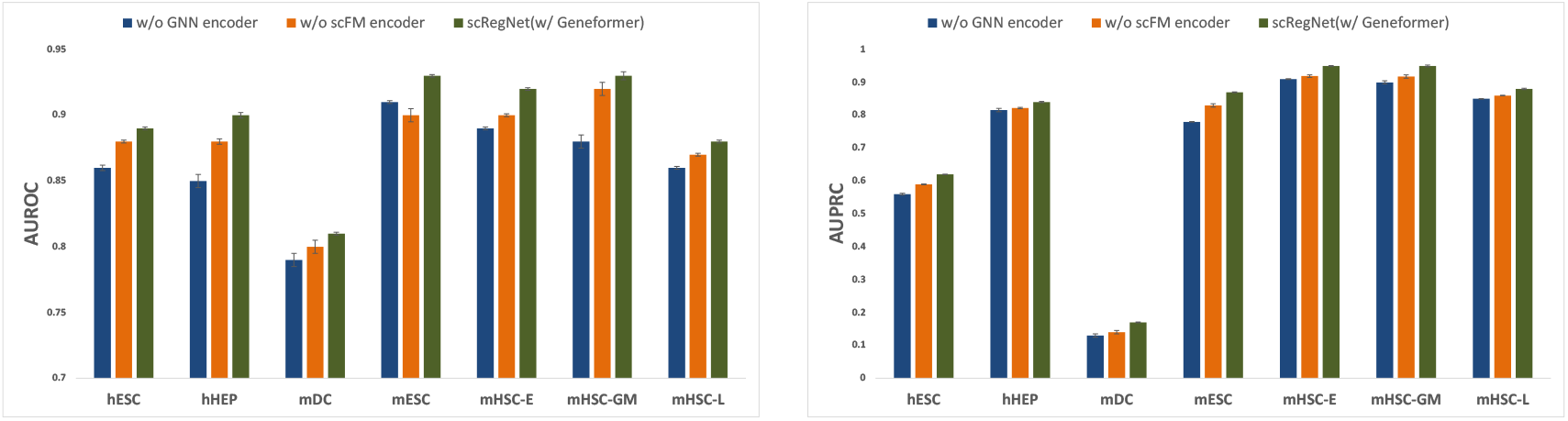
Ablation study validating the contributions of the GNN encoder and scFM (w/ Geneformer) encoder in scRegNet, evaluated using cell-type-specific GRNs. The analysis considers networks with TFs + 500 and TFs + 1000 genes, and the reported scores represent the average AUROC (left) and AUPRC (right) across both configurations, highlighting the impact of each component on model performance.

### 4.3. Impact of GNN architecture

To understand the effective choice of GNN architecture, we employ three distinct GNN architectures: GCN (Kipf and Welling, 2017), GraphSAGE (Hamilton et al., 2017), and GAT (Veličković et al., 2018). *GCN* is designed to capture the local structure of the graph by aggregating features from a node’s immediate neighbors. *GraphSAGE* builds upon GCN by enabling the aggregation of information from sampled neighborhoods, rather than requiring all neighbors to be included. *GAT* introduces attention mechanisms that weigh the importance of different neighbors during the message-passing process.

We evaluated scRegNet(w/ Geneformer) with all three of the above-mentioned GNN architectures. The evaluation results are detailed in Supplementary Table S3. This analysis reveals an interesting phenomenon: after extensive hyperparameter tuning, all GNN variants show similar performance, with only slight differences observed among them. This similarity in performance can be attributed to the sparsity of the networks (Bajaj et al., 2024). This phenomenon underscores the importance of considering graph sparsity when designing and applying GNNs. It suggests that in some cases, simpler GNN architectures may be sufficient for sparse biological networks, and that efforts to improve performance might be better directed towards graph construction and feature engineering rather than increasing model complexity.

### 4.4. Robustness Study

While our methodology leverages experimentally validated gene regulations, we acknowledge that in practice, these interactions may contain noise and false positives. Therefore, it is essential to evaluate the robustness of our model when subjected to noisy priors. To address this, we assessed the performance of scRegNet(w/ Geneformer), under various levels of noise-corrupted training data. We introduced controlled perturbations to the priors by flipping the labels of positive instances to negative and vice versa, simulating noise levels of 1%, 2%, 3%, and 4% in the training data. For each noise level, we generated 10 distinct noise-corrupted priors to ensure diverse variations. The performance of scRegNet(w/ Geneformer) was evaluated against each corrupted prior, with results visualized through box plots to illustrate performance variations.

To contextualize the robustness of scRegNet(w/ Geneformer), we benchmarked its performance against the baseline method, GENELink. The results demonstrated the stability and resilience of scRegNet under noisy training data, with consistently superior performance compared to GENELink, even as noise levels increased. These findings, depicted in Figure 3, underscore the reliability of scRegNet(w/ Geneformer) in leveraging experimentally validated gene regulations, even in the presence of noise. This robustness positions the model as a reliable choice for real-world applications with noisy training data.

**Fig. 3:**
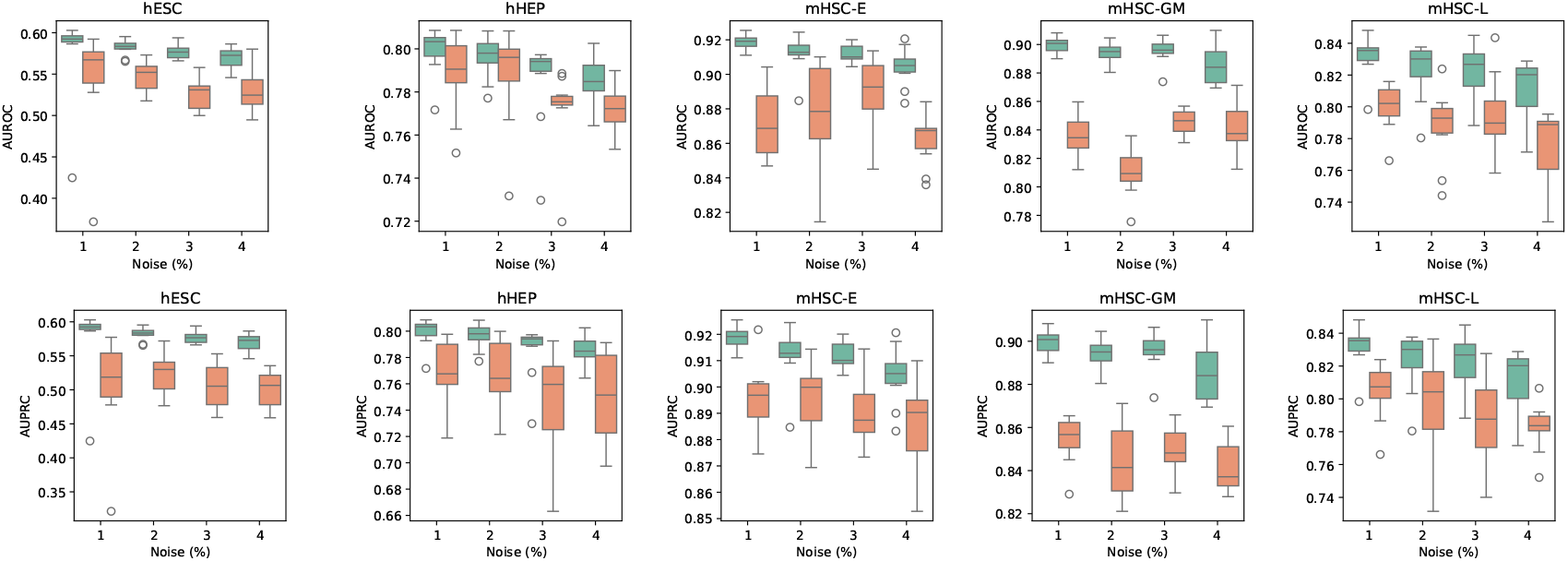
Performance comparison of scRegNet-Geneformer(green) vs. GENELink(orange) under increasing noise levels in cell-type-specific GRNs. The evaluation was conducted on networks containing TFs + 500 genes, with noise in the training dataset incrementally increased from 1% to 5%. Box plots illustrate the robustness of scRegNet in comparison to GENELink as noise levels rise, highlighting the model’s stability across varying perturbations.

## 5. Conclusions and Discussions

In this study, we introduced scRegNet, a novel and effective framework for gene regulatory link prediction by combining scFMs with GNNs. Our comprehensive evaluations across seven scRNA-seq benchmark datasets demonstrate that scRegNet consistently outperforms traditional and graph-only approaches, achieving superior AUROC and AUPRC scores. The integration of single-cell foundation models, such as scBERT, Geneformer, and scFoundation, has proven pivotal in improving the accuracy of gene regulatory link prediction. This underscores the potential of leveraging large-scale, pre-trained models that encapsulate rich biological context through self-supervised learning from extensive scRNA-seq datasets.

While our current approach demonstrates promising results, it still has several limitations. One notable challenge lies in the reliance of ground-truth TF-DNA interactions in the training data, which may not always be available for new cell types. To address this, future work will focus on incorporating the graph topology knowledge of existing TF-DNA interactions into the foundation model pre-training phase and directly do a GRN inference on the scRNA-seq data only for new cell types. This enhancement will enable broader applicability across diverse downstream tasks. Additionally, the integration of foundation models and GNNs in scRegNet remains relatively shallow and non-interactive, as each modality is currently encoded independently and fused only at the prediction stage. To overcome this limitation, we propose exploring more sophisticated and integrated fusion strategies, such as cross-modal attention mechanisms (Tsai et al., 2019), where foundation model embeddings and graph embeddings mutually inform one another through iterative attention-based updates.

## 7. Acknowledgments

Our work is sponsored by the NSF NAIRR Pilot with PSC Neocortex and NCSA Delta, Commonwealth Cyber Initiative, Children’s National Hospital, Fralin Biomedical Research Institute (Virginia Tech), Sanghani Center for AI and Data Analytics (Virginia Tech), Virginia Tech Innovation Campus, and generous gifts from Cisco Research and the Amazon + Virginia Tech Center for Efficient and Robust Machine Learning. Yizhi Wang and Yue Wang are supported by National Institutes of Health under Grants HL111362, CA271891, MH110504, and NS123719.

