## Supplementary Material for "Prediction of Gene Regulatory Connections with Joint Single-Cell Foundation Models and Graph-Based Learning"

### S1. Single-Cell Foundation Models (scFMs)

scFMs are trained on broad single-cell data using large-scale self-supervision, allowing them to be adapted for a wide range of downstream tasks. These models exhibit notable differences in training size, model architecture, design parameters, input data, and other technical features, showcasing their distinct strategies for managing single-cell transcriptomic information.. Here, we provide a brief overview of the scFMs utilized in our study: Geneformer, scFoundation, and scBERT. It's important to note that Geneformer includes additional pre-trained models with varying training sizes and design configurations; however, we have employed the largest pre-trained model available to date for this analysis.

#### S1.1. scBERT

scBERT follows bidirectional encoder representations from transformers (BERT)-based approach for pre-training and is trained using a masked learning objective – masked gene expression values are predicted as a function of the other gene embeddings in the cell. scBERT employs a matrix decomposition variant of the Transformer, known as Performer, to handle longer sequence lengths efficiently. With $L = 6$ layers, scBERT outputs an embedding size of 200 for each gene. The model is pre-trained via imputation on 5 million cells belonging to a variety of cell types from different sources. To generate emeddings for scBERT, we first requested the checkpoint and data from the corresponding authors. The environment was set up using the scBERT GitHub repository. Log-normalization was performed and cells with less than 200 expressed genes were filtered out.

#### S1.2. scFoundation

scFoundation is pre-trained over 50 million single-cells sourced from a wide range of organs and tissues originating from both healthy and donors with a variety of diseases and cancer types It employs xTrimogene as a backbone model, a scalable transformer-based architecture that includes an embedding module and an asymmetric encoder-decoder. The encoder is designed to only process nonzero and non-masked gene expression embeddings from the input matrix $\mathbf{X}$, reducing computational load and thus enabling the application of "vanilla transformer blocks to capture gene dependency without any kernel of low-rank approximation". The encoder input is formed as:

$$I = Autobin(X \odot M_{nonzero}) + Lookup(genes) \tag{8}$$

where $\odot$ denotes the element-wise product, $M_{\text{nonzero}}$ is a mask identifying non-zero elements, and Autobin converts expression values into discretized tokens. The encoder generates gene-level representations using multi-head self-attention:

$$I_{encoder} = Transformer(f_Q(I), f_K(I), f_V(I)) \tag{9}$$

where $f_Q$, $f_K$, and $f_V$ are linear projections for query, key, and value. These encoded embeddings are then recombined with the zero-expressed gene embeddings at the decoder stage to reconstruct transcriptome-wide embedded representations.

$$I_{full} = W_p(I_{encoder} \oplus I_{zero} \oplus I_{masked}) + b_p \tag{10}$$

where $\oplus$ represents concatenation, and $W_p$, $b_p$ are learned parameters that project the decoder's embedding size. The final gene-level representations are then produced by the decoder through:

$$I_{decoder} = Transformer(f_Q(I_{full}), f_K(I_{full}), f_V(I_{full})) \tag{11}$$

scFoundation is pre-trained using read-depth-aware (RDA) modeling, an extension of masked language modeling developed to take the high variance in read depth of the data into account. The raw gene expression values are pre-processed using hierarchical Bayesian downsampling in order to generate the input vectors, which can either be the unchanged gene expression profile or where downsampling has resulted in a variant of the data with lower total gene expression counts. After gene expression has been normalized, raw and input gene expression count indicators are represented as tokens which are concatenated with the model input, allowing the model to learn relationships between cells with different read depths.

To generate scFoundation embeddings, we initialized the scFoundation class shared at the official scFoundation GitHub repository. The *01B-resolution* pre-trained model checkpoint was loaded and the embeddings were generated while setting the $input\_type = singlecell$ and $tgthighres = f2$.

#### S1.3. Geneformer

Geneformer employs 20 transformer units, each consisting of a self-attention layer and an MLP layer. The model is pre-trained on Genecorpus-95M, which comprises approximately 95 million human single-cell transcriptomes from a broad range of tissues obtained from publicly available data. Before feeding the data into the model, gene expression values are converted into rank value encodings. This method provides a non-parametric representation of each single-cell transcriptome by ranking genes based on their expression levels in each cell and normalizing these ranks within the entire dataset. Consequently, housekeeping genes, which are ubiquitously highly expressed, are normalized to lower ranks, reducing their influence. Rank value encodings for each single-cell transcriptome are then tokenized, allowing genes to be stored as ranked tokens instead of their exact transcript values. Only genes detected within each cell are stored, thus reducing the sparsity of the data. When input into the model, genes from each single-cell transcriptome are embedded into a 896-dimensional space. The model is pre-trained using a masked learning objective, masking a portion of the genes and predicting the masked genes, which is intended to allow the model to learn gene network dynamics.

To generate embeddings for Geneformer, we downloaded the repository, including pre-trained model checkpoints, from Hugging Face. We pre-processed the raw expression files to ensure the correct naming of columns and then fed them into the Geneformer tokenizer (TranscriptomeTokenizer). Once the dataset had been tokenized, we extracted embeddings using the pre-trained checkpoint (20-layer model) with the *EmbExtractor* method.

**Table S1.** Comparision of the large-scale single-cell foundation models used in our study

| | **Geneformer** | **scFoundation** | **scBERT** |
|---|---|---|---|
| **Training Size: #single-cells** | 95M | 50M | 1M |
| **Model architecture** | Encoder Only | Asymmetric Encoder-Decoder | Encoder Only |
| **Design (Layer-Head-Dim)** | Transformer: 20-14-896 | Encoder Transformer: 12-12-768<br>Decoder Performer: 6-8-512 | Performer: 6-10-200 |
| **Input value** | Ranked normalized expression values | continuous normalized expression values | binned normalized expression values |
| **Number of input genes** | 4096 genes with different ranks | 19,264 protein-coding or mitochondrial genes | 16,906 genes |
| **Masking** | Non-Zero genes only | Zero and Non-Zero | Non-Zero genes only |
| **Pre-training Date** | Apr-24 | Jun-24 | Dec-21 |

### S2. Baseline Methods

We introduce the baseline methods utilized in our study, which encompass a diverse set of approaches for gene regulatory link prediction. These methods include traditional statistical techniques, machine learning algorithms, and deep learning models, applied to single-cell RNA-seq.

- GNNLink (Mao et al., 2023) is a graph neural network model that uses a GCN-based interaction graph encoder to capture gene expression patterns.
- GENELink (Chen and Liu, 2022) proposes a graph attention network (GAT) approach to infer potential GRNs by leveraging the graph structure of gene regulatory interactions.
- GNE (gene network embedding) (Kc et al., 2019) proposes a multilayer perceptron (MLP) approach to encode both gene expression profiles and network topology for predicting gene dependencies.
- CNNC (Yuan and Bar-Joseph, 2019) proposes inferring GRNs using deep convolutional neural networks (CNNs).
- DeepDRIM (Chen et al., 2021) is a supervised deep neural network that utilizes images representing the expression distribution of joint gene pairs as input for binary classification of regulatory relationships, considering both target TF-gene pairs and potential neighbor genes.
- GRN-transformer (Shu et al., 2022) is a weakly supervised learning method that utilizes axial transformers to infer cell type-specific GRNs from single-cell RNA-seq data and generic GRNs.
- Pearson correlation coefficient (PCC) (Salleh et al., 2015) is a traditional statistical method for measuring the linear correlation between two variables, often used as a baseline for GRN inference.
- GRNBoost2 (Moerman et al., 2019) is a gradient boosting-based method for GRN inference.
- GENIE3 (Huynh-Thu et al., 2010) is a random forest-based machine learning method that constructs GRNs based on regression weight coefficients, and won the DREAM5 In Silico Network Challenge in 2010.

### S3. Hyperparameter Search Space

For each combination of dataset and model, we conducted 50 optimization trials, ensuring a thorough investigation of potential configurations. Optuna's Tree-structured Parzen Estimator (TPE) was employed to intelligently navigate the search space, prioritizing regions with higher potential based on previous trials. We also applied early stopping criteria to save computational resources by halting underperforming trials early.

**Table S2.** * Applicable only for Graph Attention Network models

| hyperparameter | search space | type |
|---|---|---|
| learning rate | [1e-5, 1e-2] | continual |
| weight decay | [1e-5, 1e-4] | continual |
| dropout | [0.1, 0.8] | continual |
| #GNN layers | [1, 6] | discrete |
| #MLP layers | [1, 6] | discrete |
| batch size | [32, 56] | discrete |
| hidden dimension | [4, 256] | discrete |
| #attention heads* | [1, 8] | discrete |
| reduction* | [concatenate, mean] | discrete |
| alpha* | [0.01, 0.5] | continual |
| optimizer | [Adam, RMSprop, SGD] | discrete |

### S4. End-to-End Training Algorithm

Algorithm S1 details the end-to-end training process of scRegNet for gene regulatory link prediction using a combination of a pre-trained single-cell foundation model and graph neural network (GNN).

**Algorithm S1** End-to-End Training for Gene Regulatory Link Prediction

**Input**: Pre-trained Single-Cell Foundation Model $F$, raw count matrix $\mathbf{X}$,
dataset $\mathcal{D}$ with positive and negative gene pairs, Gene Set $\mathcal{T}$
**Output**: Optimized Model for gene regulatory link prediction

**Dataset Preparation**: Split dataset $\mathcal{D}$ into training set $\mathcal{D}_{\text{train}}$, validation set $\mathcal{D}_{\text{val}}$, and test set $\mathcal{D}_{\text{test}}$
**Embedding Extraction**: Compute gene embeddings using $F$: $\mathbf{E} = F(\mathbf{X})$
**Attentive/Avg Pooling**: $p_t = POOLING([E_t^{cell1}, E_t^{cell2}, ..., E_t^{celln}]), \quad \forall t \in \mathcal{T}$
**Model Freezing**: Freeze the layers of the pre-trained foundation model $F$
**GNN Initialization**: Initialize node embeddings for the GNN: $v_t^0 \leftarrow x_t$, $\forall t \in \mathcal{T}$
**for** each training epoch **do**
  **Graph Encoding**: $\hat{v}_t \leftarrow \text{GNN}(v_t^0, \mathcal{D}_{\text{train}})$
  **for** each batch $b$ containing $(\text{TF}, \text{target})$ gene pairs in $\mathcal{D}_{\text{train}}$ **do**
    **Joint Embedding Construction**: Concatenate transformer and graph embeddings:

$$Z_{\text{joint}}[\text{TF}_b] = \text{CONCAT}(\mathbf{p}_{\text{TF}_b}, v_{\text{TF}_b}), \quad Z_{\text{joint}}[\text{target}_b] = \text{CONCAT}(\mathbf{p}_{\text{target}_b}, v_{\text{target}_b})$$

    **MLP Processing**: $e_{\text{TF}} = \text{MLP}_1(Z_{\text{joint}}[\text{TF}]), \quad e_{\text{target}} = \text{MLP}_2(Z_{\text{joint}}[\text{target}])$
    **Link Prediction**: $h_b = \text{CONCAT}(e_{\text{TF}_b}, e_{\text{target}_b}), \quad \text{prob} = \text{softmax}(\mathbf{W}h_b + \mathbf{b})$
    **Loss Calculation**: Compute binary cross-entropy loss $\mathcal{L}_{\text{CE}}$
    **Parameter Update**: Optimize the GNN and MLP weights via backpropagation
  **end for**
**end for**
**Evaluation**: Assess the model on $\mathcal{D}_{\text{test}}$ and report performance metrics (AUROC, AUPRC)
**return** Optimized Model $\mathcal{M}^*$

### S5. Additional Experimental Results

Comparative analysis of link prediction performance in GRNs using different GNN architectures as backbone encoders for scRegNet, with Geneformer serving as the foundation model. The experiments were conducted using two different sets of most variable genes combined with transcription factors (TFs): (a) 500 most variable genes and (b) 1000 most variable genes.

**Table S3.** Comparative Analysis of Link Prediction Performance in GRNs Using Popular GNN Variants as Backbone Graph-Based Encoders for scRegNet, with Geneformer Serving as the Foundation Model Backbone.

| Cell Type | | GCN | SAGE | GAT |
|---|---|---|---|---|
| hESC | AUROC | 0.886 ± 0.002 | 0.884 ± 0.004 | 0.886 ± 0.004 |
| | AUPRC | 0.622 ± 0.004 | 0.623 ± 0.003 | 0.614 ± 0.002 |
| hHEP | AUROC | 0.902 ± 0.002 | 0.901 ± 0.003 | 0.901 ± 0.003 |
| | AUPRC | 0.841 ± 0.003 | 0.839 ± 0.004 | 0.838 ± 0.003 |
| mDC | AUROC | 0.808 ± 0.003 | 0.788 ± 0.001 | 0.808 ± 0.002 |
| | AUPRC | 0.167 ± 0.002 | 0.158 ± 0.002 | 0.167 ± 0.001 |
| mESC | AUROC | 0.926 ± 0.001 | 0.926 ± 0.004 | 0.926 ± 0.002 |
| | AUPRC | 0.860 ± 0.001 | 0.859 ± 0.001 | 0.859 ± 0.003 |
| mHSC-E | AUROC | 0.923 ± 0.001 | 0.923 ± 0.001 | 0.921 ± 0.004 |
| | AUPRC | 0.941 ± 0.002 | 0.941 ± 0.003 | 0.939 ± 0.002 |
| mHSC-GM | AUROC | 0.931 ± 0.002 | 0.930 ± 0.002 | 0.929 ± 0.003 |
| | AUPRC | 0.939 ± 0.001 | 0.939 ± 0.001 | 0.938 ± 0.003 |
| mHSC-L | AUROC | 0.879 ± 0.002 | 0.877 ± 0.002 | 0.878 ± 0.002 |
| | AUPRC | 0.879 ± 0.002 | 0.874 ± 0.001 | 0.876 ± 0.003 |

(a) TFs + 500 Most Variable Genes

| Cell Type | | GCN | SAGE | GAT |
|---|---|---|---|---|
| hESC | AUROC | 0.883 ± 0.002 | 0.882 ± 0.002 | 0.885 ± 0.002 |
| | AUPRC | 0.624 ± 0.003 | 0.620 ± 0.004 | 0.625 ± 0.001 |
| hHEP | AUROC | 0.905 ± 0.004 | 0.905 ± 0.003 | 0.905 ± 0.002 |
| | AUPRC | 0.844 ± 0.001 | 0.844 ± 0.002 | 0.843 ± 0.002 |
| mDC | AUROC | 0.838 ± 0.002 | 0.835 ± 0.003 | 0.838 ± 0.004 |
| | AUPRC | 0.162 ± 0.002 | 0.146 ± 0.002 | 0.162 ± 0.003 |
| mESC | AUROC | 0.931 ± 0.002 | 0.931 ± 0.001 | 0.930 ± 0.003 |
| | AUPRC | 0.866 ± 0.002 | 0.866 ± 0.002 | 0.865 ± 0.004 |
| mHSC-E | AUROC | 0.941 ± 0.004 | 0.941 ± 0.002 | 0.940 ± 0.002 |
| | AUPRC | 0.956 ± 0.003 | 0.954 ± 0.001 | 0.953 ± 0.002 |
| mHSC-GM | AUROC | 0.936 ± 0.003 | 0.937 ± 0.004 | 0.935 ± 0.003 |
| | AUPRC | 0.949 ± 0.002 | 0.950 ± 0.001 | 0.948 ± 0.003 |
| mHSC-L | AUROC | 0.876 ± 0.001 | 0.876 ± 0.003 | 0.876 ± 0.002 |
| | AUPRC | 0.865 ± 0.001 | 0.865 ± 0.001 | 0.864 ± 0.004 |

(b) TFs + 1000 Most Variable Genes
